# Sexually dimorphic effects of Amylin 1 receptor activation in trigeminovascular neurons

**DOI:** 10.1101/2024.01.12.575235

**Authors:** Alejandro Labastida-Ramírez, Eloisa Rubio-Beltran, Philip R. Holland, Jan Hoffmann

## Abstract

**Background:** Migraine is more prevalent in women, and although the mechanisms involved in this disparity remain poorly understood, an interaction between the trigeminovascular system and cycling estrogen levels in biologically-predisposed women has been suggested. We investigated the role of amylin 1 (AMY_1_) receptor activation in the modulation of the trigeminal nociceptive system in female rats across the estrous cycle in cycle stages with falling and rising estrogen levels and compared these to the responses in males.

**Methods:** We recorded neuronal activity *in vivo* within the trigeminocervical complex (TCC) and examined the effects of targeting AMY_1_ receptors on ongoing spontaneous and dural stimulus-evoked firing rates of trigeminovascular neurons. The selective AMY_1_ receptor agonist pramlintide and AMY_1_ receptor antagonist AC 187 were used. Estrous cycle stages were identified via cytology from vaginal smears.

**Results:** Administration of pramlintide increased the spontaneous activity and dural stimulus-evoked neuronal responses in the TCC, only during falling estrogen phases of the female estrous cycle. Moreover, the administration *per se* of AC 187 decreased spontaneous evoked firing rates of central trigeminovascular neurons in females and males, whereas pretreatment with AC 187 prevented pramlintide-induced increases in spontaneous activity and dural stimulus-evoked responses in females with falling estrogen levels.

**Conclusion:** AMY_1_ receptors modulate the trigeminal nociceptive system. The facilitating effect is most pronounced in female rats during falling estrogen phases of the estrous cycle. Our data also supports selective AMY_1_ receptor antagonists as potentially effective targets for the treatment of migraine.

## Background

Migraine is the leading cause of disability in young women, affecting them during productive years of their adulthood [1, 2]. Unfortunately, a remarkable increase in migraine prevalence, three times more common than in men, occurs in women after menarche, with hormonal fluctuations influencing migraine attack susceptibility during different reproductive milestones, such as menstruation and ovulation, as well as pregnancy and lactation [3, 4]. Moreover, migraine frequency is influenced by exogenous and endogenous dysfunction of the hypothalamo-pituitary-ovarian system, such as usage of hormonal contraceptives and disorders like endometriosis and polycystic ovary syndrome, respectively [3, 4]. Although the precise neurobiological mechanisms underlying this female disparity are still poorly understood, an interaction between cycling estrogen levels and the trigeminovascular system in biologically-predisposed women has been suggested [5, 6].

Calcitonin gene-related peptide (CGRP) is a key neuropeptide involved in migraine pathophysiology, which is widely expressed throughout the nociceptive trigeminovascular system [7, 8]. Higher levels of this peptide have been measured in different biological fluids of migraine patients during migraine attacks [9]. Therefore, drugs targeting CGRP or the canonical CGRP receptor have emerged as breakthrough therapies in migraine treatment. Interestingly, different studies have reported sexual dimorphism in the plasma levels of CGRP, as well as in the expression of the components of its canonical receptor [3, 10]. Moreover, the expression of sex hormone receptors in the peptidergic trigeminovascular system indicates that trigeminal neurons may respond to variations in the levels of these hormones, suggesting a complex crosstalk between fluctuating sex hormones and CGRP signaling [3].

In addition to the canonical CGRP receptor, recent studies have shown that CGRP can also activate a second CGRP responsive receptor, the amylin 1 (AMY_1_) receptor, which is also potently activated by the brain-gut peptide amylin [9, 11]. AMY_1_ receptors are also expressed in trigeminal fibers and have recently been linked to migraine pathophysiology [12]. In contrast to the CGRP receptor, the AMY_1_ receptor undergoes scarce internalization after repeated agonist stimulation [13], suggesting this receptor will remain accessible to therapies during migraine attacks. Furthermore, amylin signaling in the brain is highly dependent on sex hormones, diminishing by ovariectomy and recovering by estradiol replacement [14]. This suggests a potential crosstalk of amylin signaling and sex hormones in migraine pathophysiology. However, the influence of sex hormones on AMY_1_ receptor signaling in the trigeminovascular system remains to be determined. Therefore, the aim of the present study was to investigate the effect of AMY_1_ receptor activation in the modulation of trigeminal nociceptive transmission during falling and rising estrogen levels in a validated clinically relevant rodent model of migraine-related pain processing.

## Methods

All animal care and experimental procedures complied with the UK Home Office Animals (Scientific Procedures) Act 1986 and were approved by King’s College London, Animal Welfare and Review Body. Animal studies are reported in compliance with ARRIVE 2.0 guidelines [15].

### Experimental design

Adult male and female Sprague Dawley (200-350 gr; Envigo UK Ltd) rats were used throughout. They were group-housed in the animal facility under standard conditions in temperature and light controlled rooms for at least 7 days before use, with access to water and food *ad libitum*. All animals were randomly assigned into the different experimental protocols, females were then pseudo randomized based on estrous cycle stages (see next section). In each animal, only one experiment was conducted, and all analyses were performed by an observer blinded to the experimental grouping and stage of the estrous cycle. At the end of each experiment, all animals were humanely euthanized with an intravenous overdose of 200 mg/kg pentobarbitone (Euthatal, Merial, UK).

### Vaginal smear and surgical preparation

The stages of the rodent estrous cycle were identified through microscopic evaluation of the cell composition present in air dried Cresyl violet-stained (25 µl) vaginal smears. Two vaginal smears were taken on the day of the experiment (9:30 am and 17:30 pm) and matched to the morning vaginal and rectal temperature of the rat [16, 17]. As previously described, this allows categorizing the rodent estrous cycle into different phases: proestrus, estrus, metestrus, diestrus I and diestrus II [16, 17]. In order to maintain blinding, the vaginal smears and staining were not conducted by the person conducting the experiment.

The surgical procedure, physiological monitoring and electrophysiological recording methods have been reported previously [18, 19]. In brief, rats were initially anesthetized with isoflurane (IsoFlo 5%, Zoetis, UK) and maintained with an intravenous propofol infusion (PropoFlo Plus, 30-50 mg/kg/hr, Zoetis, UK). The depth of anesthesia was judged by a negative paw withdrawal test, mean arterial pressure levels, the absence of ocular reflexes, and changes to expired CO_2_. The femoral veins and artery were cannulated for intravenous administration of drugs and for continuous monitoring of mean arterial pressure, respectively. Subsequently, the rats were placed in a stereotaxic frame, and continuously monitored for mean arterial blood pressure and body temperature. Animals were mechanically ventilated with O_2_-enriched air, and end expired CO_2_ levels maintained between 3.5% and 4.5% [18, 19].

### In vivo electrophysiological recordings

The parietal bone was drilled to provide access to the cranial dura mater overlying the middle meningeal artery where a bipolar simulating electrode was placed to stimulate trigeminal perivascular sensory afferents (8-15 V, 0.5 Hz, 0.3-0.5 ms, 20 square wave electrical pulses) [19]. An ipsilateral C1 hemilaminectomy was performed and the dura mater was pierced to expose the brainstem at the level of the caudal medulla [18, 19]. A tungsten recording electrode (0.5-1 MΩ) was lowered into the trigeminocervical complex (TCC) region of the brainstem at 5-µm with a motorized manipulator (Scientifica, UK). The neuronal signal was amplified, filtered, and fed to a gated amplitude discriminator and an analogue-to-digital converter and to a microprocessor-based computer for analysis using Spike 2 v8.12. After completion of the surgery, animals were rested for at least 30 min before recording commenced.

Extracellular recordings were made from neurons in the TCC and identified as having cutaneous facial receptive fields in the V1 dermatome and activated by electrical stimulation of the dural afferents. Trains of 20 stimuli were delivered at 5-min intervals to assess the baseline response to dural electrical stimulation. Stimulation parameters were optimized to ensure that stimulation activated both Aδ-fiber (3-20 ms latency range) and C-fiber (20-80 ms latency range) neurons [18, 19]. Responses were analyzed using post-stimulus histograms with a bin width of 1 ms and sweep duration of 100 ms. Innocuous cutaneous responses were assessed by brushing the receptive field with a cotton tip applicator. Ongoing spontaneous activity (spikes/second) was recorded throughout and measured for analysis taken for 30 sec preceding the dural stimulation. Peri- and post-stimulus time histograms of neural activity were displaced and analyzed with Spike 2 v8.12. Once stable baseline values of the stimulus-evoked responses were achieved (average of three stimulation series), responses were tested for 90 min following drug administration. To investigate the involvement of AMY_1_ receptors on basal trigeminocervical neuronal responses, a selective agonist or antagonist were administered intravenously. In a subset of experiments, the antagonist was administered intravenously 5 minutes before agonist injection (see below).

### Compounds

Apart from the anesthetics, the compounds used in the present study (obtained from Tocris Bioscience, UK) were the selective AMY_1_ receptor agonist pramlintide (pEC_50_ 9.4), and the selective AMY_1_ receptor antagonist AC 187 (p*K*_B_ 8.0) [20, 21]. All compounds were dissolved in physiological saline, aliquoted, and frozen until the day of experiment. The intravenous doses of pramlintide (1.6 nmol/kg) and AC-187 (2 nmol/kg) are based on previous studies [22, 23]. Saline (1 ml/kg) was the vehicle control throughout as it has no significant effects in electrophysiological recordings as previously reported by our group.

### Data presentation and statistical analysis

The required number of animals used per group was based on published data and previous experience that typically sees differences in means of 28-30% (SD = 15-20%), with a two-sided alpha of 0.05 and power of 80%, which requires sample sizes of 6-8 animals [22]. All data in the text and figures are expressed as mean ± standard error of the mean (SEM). All statistical analyses were performed on GraphPad Prism 10.0.3 software. For multiple time points, an analysis of variance for repeated measures with Dunnett post hoc correction for multiple comparisons was used to measure the time course of significant drug intervention, using a 95% confidence interval. If Mauchly’s test of sphericity was violated, we made appropriate corrections to degrees of freedom according to Greenhouse-Geisser. Statistical significance was set at p < 0.05.

## Results

### General

Electrophysiological recordings were made from 78 neuronal recordings in 78 rats that responded to electrical stimulation of the cranial dura mater, and with cutaneous facial receptive fields that included predominantly the V1 dermatome. Neurons were mainly located in nociceptive specific superficial and deeper layers, laminae I-II and laminae V and VI, respectively, of the dorsal horn of the TCC at ranges of depth of 100-1000 µm.

### Effects of vehicle and pramlintide on trigeminocervical neuronal responses

As previously reported by our group, there were no significant differences between male and female responses to intravenous injection of vehicle (n = 6 each, data not shown). As there were also no significant differences to treatments (data not shown) between female responses during phases of the estrus cycle characterized by rising (proestrus, metestrus, diestrus II) or falling (estrus + diestrus I) estrogen, these data was pooled. Therefore, female data were grouped for further analysis into either falling (n = 24) or rising (n = 33) estrogen and compared to males (n = 21).

As shown in Figure 1, in males (n = 6), intravenous injection of pramlintide (1.6 nmol/kg) had no significant effect on ongoing spontaneous trigeminal neuronal firing (F_2.0, 10.1_ = 0.52, p = 0.6), responses to innocuous stimulation of the V1 cutaneous receptive field (F_2.6, 13.2_ = 0.60, p = 0.6), or stimulus-evoked responses within the Aδ-fiber (F_2.3, 11.4_ = 0.77, p = 0.5) and C-fiber (F_1.7, 8.6_ = 0.34, p = 0.7) range over the entire recording period of 90 min.

**Figure 1.**
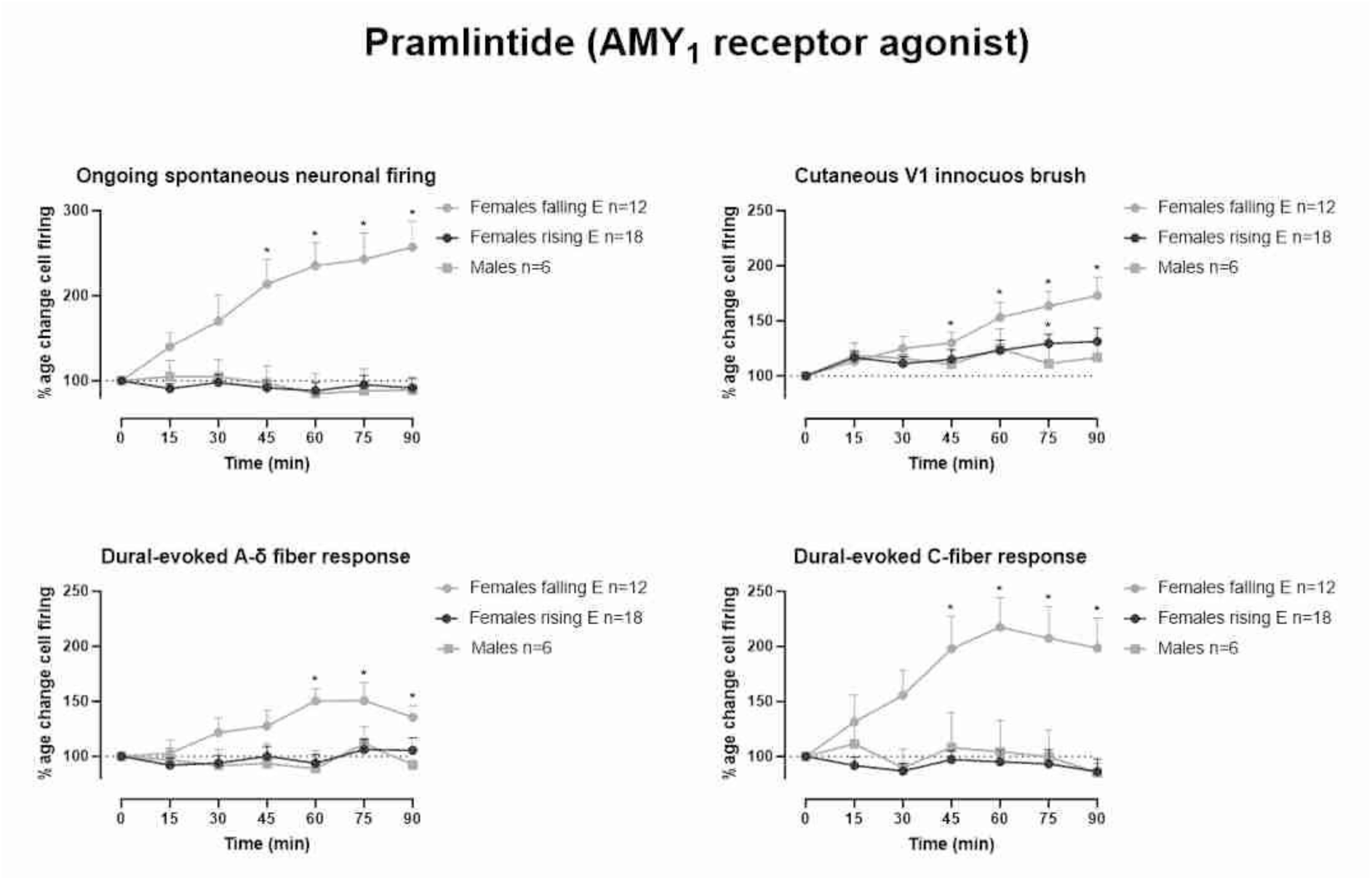
Pramlintide causes delayed activation of trigeminal neurons during phases of the female estrous cycle that are characterized by falling estrogen levels. Time course changes in (a) ongoing spontaneous trigeminocervical neuronal firing, (b) innocuous somatosensory-evoked trigeminocervical neuronal responses, (c) intracranial dural-evoked neuronal responses with inputs in the A-δ fiber range and (d) C-fiber latency range, in response to pramlintide (1.6 nmol/kg). These data have been normalized to represent the percentage change from baseline and are expressed as mean ± SEM. *p < 0.05 compared to baseline.

In females during phases of the estrous cycle that are characterized by rising estrogen levels (n = 18), no significant effect on ongoing spontaneous trigeminal neuronal firing (F_2.8, 47_ = 0.60, p = 0.6), or stimulus-evoked responses within the Aδ-fiber (F_2.6, 45_ = 1.01, p = 0.4) and C-fiber (F_3.3, 57_ = 0.57, p = 0.7) range was observed. However, pramlintide significantly increased responses to innocuous stimulation of the V1 cutaneous receptive field (F_3.0, 52_ = 3.5, p = 0.02), especially after 75 minutes (129% ± 8%).

In contrast to the observations made in male rats and in female rats in cycle stages with rising estrogen levels, female rats in phases of the estrous cycle that are characterized by falling estrogen levels (n = 18), showed a significant increase in ongoing spontaneous trigeminal neuronal firing (F_3.0, 34_ = 13.3, p < 0.0001) and responses to innocuous stimulation of the V1 cutaneous receptive field (F_2.4, 26_ = 11.7, p = 0.0001) across the entire recording period of 90 min after pramlintide. Specifically after 45 min (214% ± 29% and 130% ± 10%, respectively), and maximally at 90 min (257% ± 31% and 173% ± 17%, respectively). Additionally, as shown in Figure 1, both dural stimulus-evoked responses were significantly increased after pramlintide, with Aδ-fibers (F_3.5, 39_ = 5.7, p = 0.0016) responding initially after 60 min (150% ± 11%) and maximally at 75 min (151% ± 16%), whereas C-fibers (F_6, 66_ = 5.5, p = 0.0001) responded initially after 45 min (198% ± 27%) and maximally at 60 min (218% ± 27%).

### Effects of vehicle and AC 187 on trigeminocervical neuronal responses

As shown in Figure 2, vehicle (1 ml/kg, n = 12) had no significant effects throughout the 90-min recording period on ongoing spontaneous trigeminal neuronal firing (F_2.2, 24.3_ = 0.63, p = 0.6), or stimulus-evoked responses within the Aδ-fiber (F_1.6, 18_ = 0.61, p = 0.5) and C-fiber (F_2.2,24_ = 0.32, p = 0.8) range.

**Figure 2.**
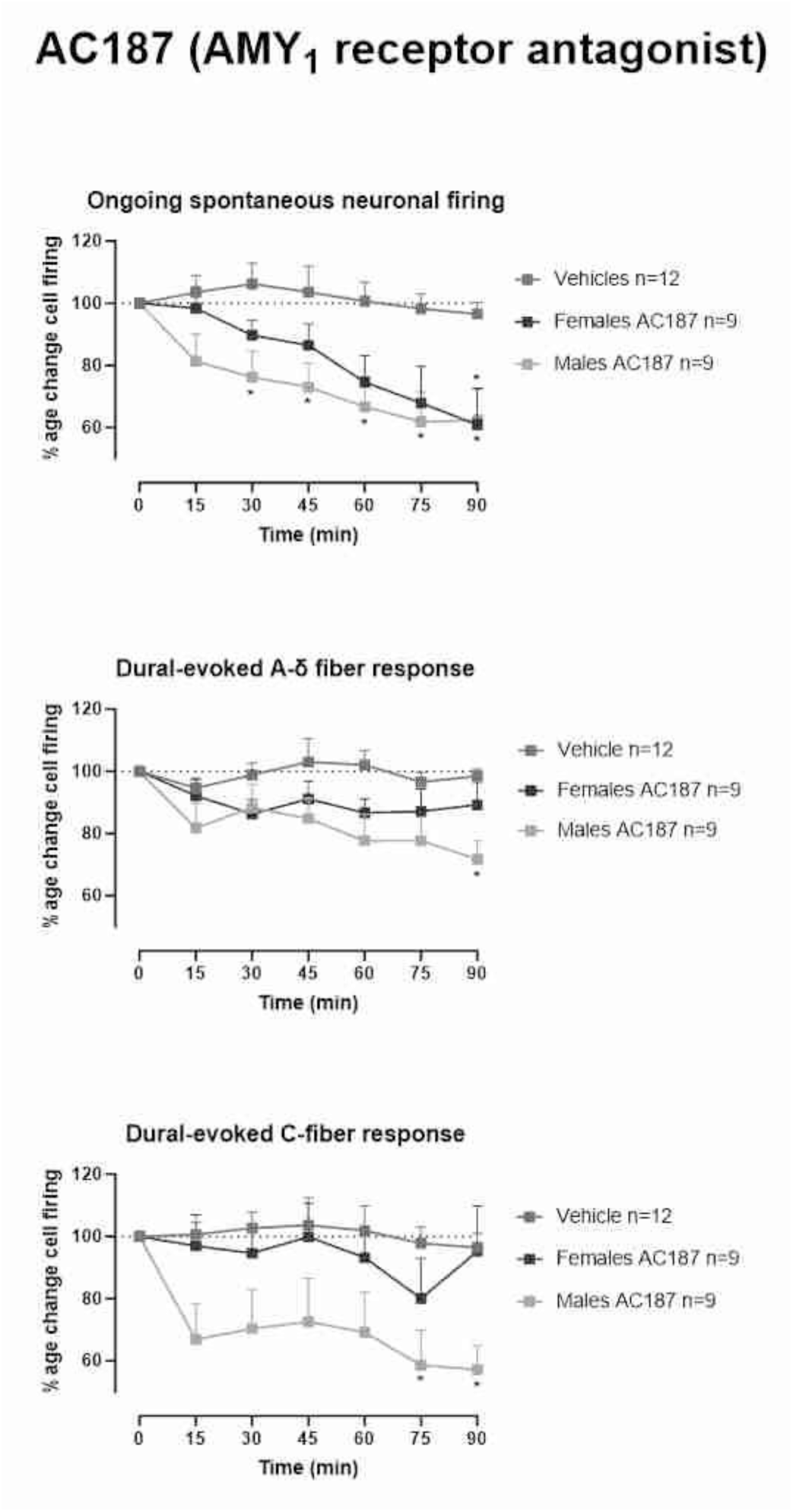
AC 187 decreases trigeminocervical neuronal responses. Time course change in (a) spontaneous trigeminocervical neuronal firing, (b) intracranial dural-evoked neuronal responses with inputs in the A-δ fiber range and (c) C-fiber latency range, in response to AC 187 (2 nmol/kg). These data have been normalized to represent the percentage change from baseline and are expressed as mean ± SEM. *p < 0.05 compared to baseline.

At 2 nmol/kg AC 187 showed in males (n = 9) a significant decrease on ongoing spontaneous trigeminal neuronal firing (F_2.7, 21.3_ = 8.9, p = 0.0007), initially after 30 min (−26% ± 7%) and maximally at 90 min (−38% ± 10%). Moreover, AC 187 significantly inhibited neuronal responses within the Aδ-fiber (F_2.9, 23.8_ = 4.3, p = 0.0151) and C-fiber range across 90 min (F_3.4, 27.6_ = 4.1, p < 0.0001), maximally at 90 min (−28% ± 6% and −43% ± 8%, respectively).

In females (n = 9, 6 rising and 3 falling estrogen phases), 2 nmol/kg AC 187 significantly inhibited ongoing spontaneous trigeminal neuronal firing (F_2.5, 20.2_ = 6.9, p = 0.003), specifically after 90 min (−39% ± 12%). Conversely, AC 187 had no significant effects in the same neuronal population responses within the Aδ-fiber (F_2.9, 23.2_ = 0.9, p = 0.5) and C-fiber (F_2.6, 20.6_ = 0.5, p = 0.6) range to dural electrical stimulation.

### Effects of AC 187 pretreatment on pramlintide-induced neuronal responses

As shown in Figure 3, in females with rising estrogen levels (n = 6), pretreatment with 2 nmol/kg AC 187 significantly inhibited pramlintide-induced spontaneous trigeminal neuronal firing (F_1.9, 9.7_ = 7.4, p = 0.0118), specifically after 45 min (−28% ± 7%), and maximally at 90 min (−43% ± 6%). Conversely, AC 187 pretreatment followed by pramlintide had no significant effects over 90 min on responses to innocuous stimulation of the V1 cutaneous receptive field (F_2.2, 11_ = 0.70, p = 0.5), or stimulus-evoked responses within the Aδ-fiber (F_3, 14.8_ = 3.05, p = 0.0619) and C-fiber (F_1.9, 9.4_ = 0.18, p = 0.8) range.

**Figure 3.**
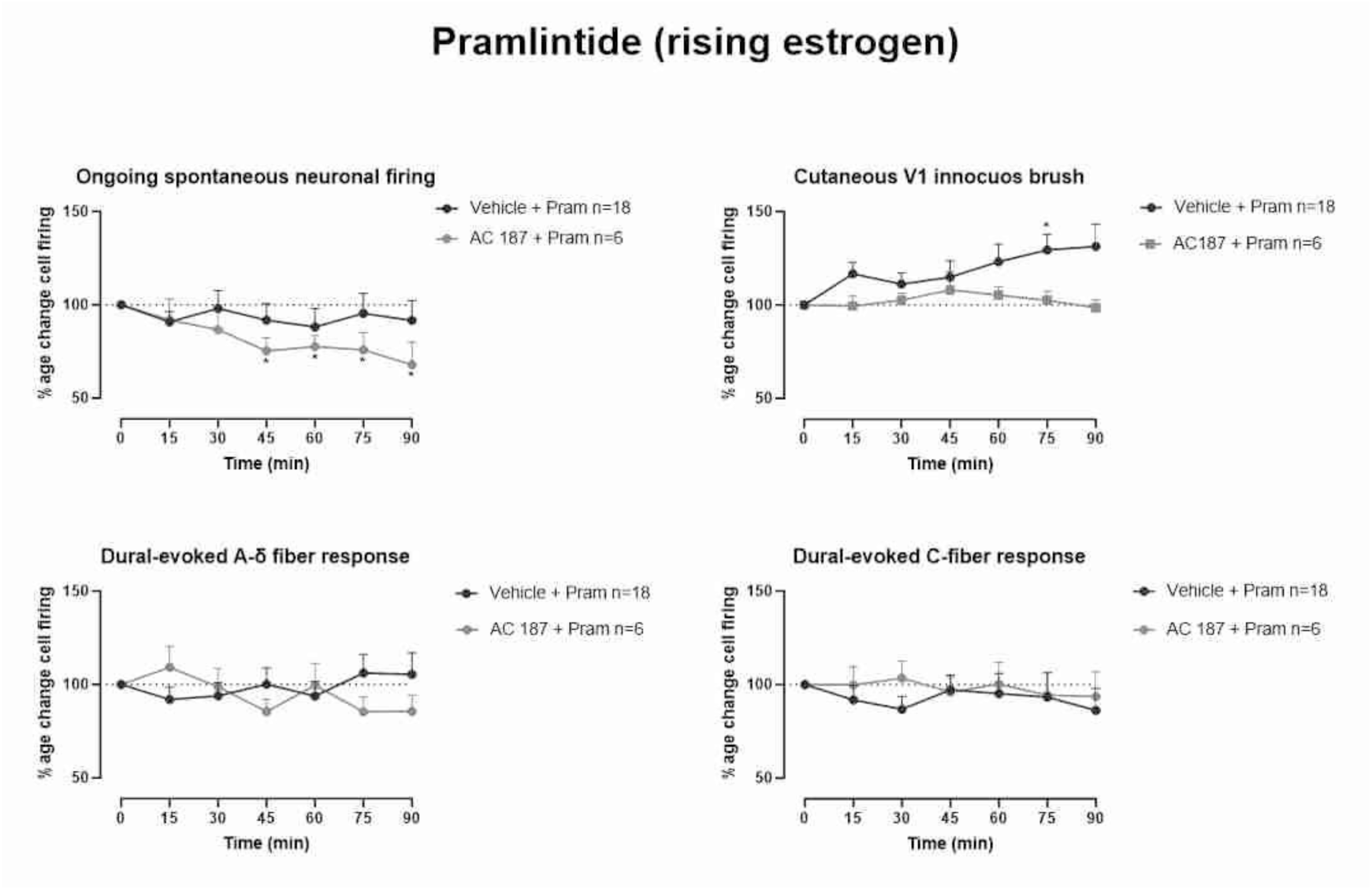
Pretreatment with AC 187 decreases trigeminocervical neuronal responses. Time course changes in (a) ongoing spontaneous trigeminocervical neuronal firing, (b) innocuous somatosensory-evoked trigeminocervical neuronal responses, (c) intracranial dural-evoked neuronal responses with inputs in the A-δ fiber range and (d) C-fiber latency range, in females with rising estrogen levels pretreated with either vehicle (1 ml/kg) or AC 187 (2 nmol/kg) followed by pramlintide (1.6 nmol/kg). These data have been normalized to represent the percentage change from baseline and are expressed as mean ± SEM. *p < 0.05 compared to baseline.

Furthermore, as shown in Figure 4, in females with falling estrogen (n = 6), pretreatment with AC 187 prevented the effects of pramlintide over 90 min on spontaneous trigeminal neuronal firing (F_1.4, 9.7_ = 3.2, p = 0.1106), or stimulus-evoked responses within the Aδ-fiber (F_2.9, 14.5_ = 3.8, p = 0.4) and C-fiber (F_1.2, 6.1_ = 3.8, p = 0.2) range. However, pretreatment with AC 187 did not prevent pramlintide increased responses to innocuous stimulation of the V1 cutaneous receptive field (F_3.1, 15.7_ = 3.8, p = 0.03), specifically after 60 min (125% ± 6%), and maximally at 90 min (129% ± 7%).

**Figure 4.**
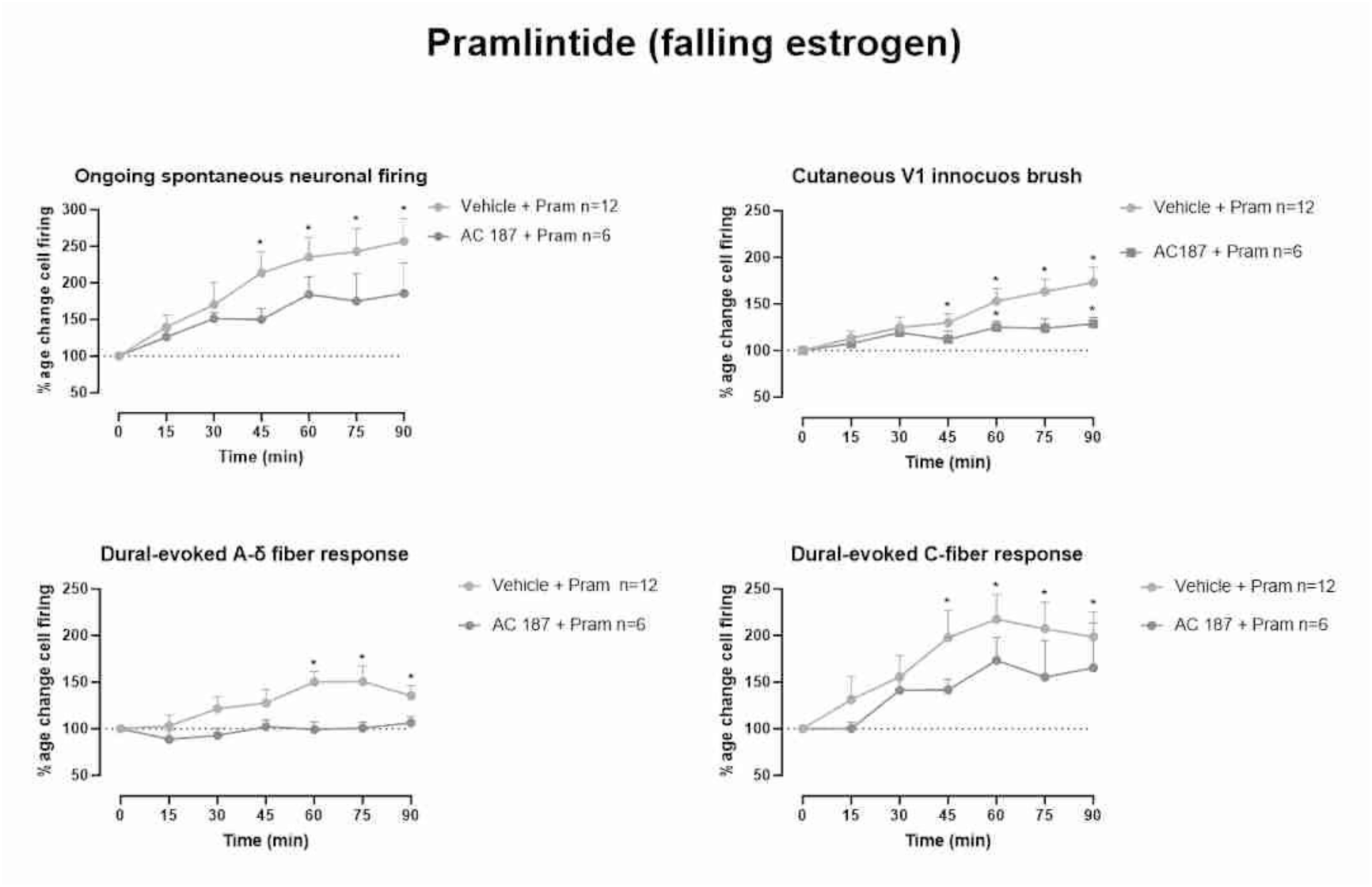
Pretreatment with AC 187 attenuates pramlintide-induced trigeminocervical neuronal responses. Time course changes in (a) ongoing spontaneous trigeminocervical neuronal firing, (b) innocuous somatosensory-evoked trigeminocervical neuronal responses, (c) intracranial dural-evoked neuronal responses with inputs in the A-δ fiber range and (d) C-fiber latency range, in females with falling estrogen levels pretreated with either vehicle (1 ml/kg) or AC 187 (2 nmol/kg) followed by pramlintide (1.6 nmol/kg). These data have been normalized to represent the percentage change from baseline and are expressed as mean ± SEM. *p < 0.05 compared to baseline.

## Discussion

In the present study, we investigated the modulation of trigeminocervical neuronal responses after pharmacological activation of AMY_1_ receptors on males and females during falling and rising estrogen levels. We demonstrated that pramlintide (1.6 nmol/kg), a selective AMY_1_ receptor agonist, causes activation and sensitization of central trigeminocervical neurons only in females, particularly during falling estrogen phases of the estrous cycle. Additionally, we investigated the potential antimigraine effect of AC 187, a selective AMY_1_ receptor antagonist, using an experimental paradigm well-validated to predict the efficacy of novel drugs [18, 24]. We established that AC 187 (2.1 nmol/kg) inhibits ongoing firing of trigeminocervical neurons in both sexes and only dural-evoked activation of trigeminocervical neurons in males. Further, antagonism of the AMY_1_ receptor with AC 187, similarly inhibited pramlintide-induced activation and sensitization of central trigeminocervical neurons.

In accordance with previous work, subcutaneous injection of low and high doses of amylin in the periorbital area (1.5 nmol and 15 nmol, respectively) did not evoke spontaneous nociceptive behaviors nor periorbital mechanical allodynia in male mice [23], although female mice were not included in the study. Similarly, in male mice, a 7-fold higher dose of intraperitoneal amylin (0.5 mg/kg) did not provoke significant migraine-like symptoms such as cutaneous hypersensitivity and light aversion, whereas female mice displayed greater sensitivity to amylin [12, 25]. Taken together, these results suggest that amylin signaling in the trigeminovascular system may contribute to the female-specific higher prevalence of migraine.

In addition to the recently described sex-dependent effects of amylin, sexually dimorphic responses to CGRP have also previously emerged in several preclinical rodent migraine models. For instance, topical and systemic administration of CGRP did not result in activation nor sensitization of meningeal nociceptors in male rats [26]. In contrast, acute responses to dural or cerebellar CGRP administration induced headache-like behavioral responses only in females [27, 28], as well as a female-specific sensitization to CGRP following peripheral and central priming [27]. Moreover, in ovariectomized females, a higher release of CGRP from perivascular trigeminal nerves was revealed after estradiol supplementation [29]. Thus, as it has become evident that female sex hormones play a role in migraine pathophysiology, with hormonal fluctuations influencing migraine attack occurrence and CGRP release (reviewed in [3]), we decided to determine the role of AMY_1_ receptor activation throughout the natural estrous cycle of the rat.

Interestingly, only females during falling estrogen phases of the estrous cycle responded and were sensitized to activation of AMY_1_ receptors by pramlintide (Figure 1). This is in line with Martin et al., who previously reported enhanced sensitization of the trigeminal system after capsaicin administration, and CGRP release, during the later halves of proestrus and estrus, which represent stages of the rodent estrous cycle during and immediately following an abrupt decline in ovarian hormones [30]. Thus, activation of AMY_1_ and CGRP receptors is a more potent promoter of trigeminal nociception in females during falling estrogen phases of the estrous cycle. However, an enhanced susceptibility to pain behaviors after serotonin and complete Freund’s adjuvant administration has also been shown in females during estrus and proestrus [31, 32]. Moreover, trigeminal mechanoreceptive neurons of females during these estrous phases exhibit lower periorbital mechanical pain thresholds and enlarged facial receptive fields [33, 34]. This indicates this is rather a naturally and cyclically occurring phenomenon, which reduces the threshold for activation of nociceptors and consequently predisposes females to experience greater pain intensities and larger areas of mechanical-induced allodynia, that seem independent of a particular neuropeptide system. Indeed, women are not only more sensitive to acute experimental pain [35], but they also report greater pain perception and sensory responses to experimental trigeminal sensitization during the menstrual phase compared to the luteal phase and males [36]. However, the relationship between pain perception and sex hormones, and the underlying mechanisms and the role of individual sex hormones in modulating nociception are complex and far from understood. In relation to migraine, experimental data suggest that estrogen influences nociceptor function as well as CGRP receptor expression and release at multiple levels of the peripheral and central trigeminovascular system [3, 35]. Further studies, beyond the scope of our study, are clearly needed to determine which estrogen receptors are involved in the modulation of AMY_1_ receptor expression and amylin release in the different components of the trigeminovascular system during the different phases of the estrous cycle.

Also, we studied the effect *per se* of a selective AMY_1_ receptor antagonist in a validated *in vivo* preclinical model of trigeminal nociception [24]. Similar to drugs that antagonize the CGRPergic system [22, 24], administration of AC 187 inhibited ongoing spontaneous firing rates and dural-evoked activation of trigeminocervical neurons in males. However, females were less sensitive to the same dose of AC 187, as only spontaneous firing rates were inhibited but neither dural-evoked activation of A-δ fibers nor C-fibers were inhibited (Figure 2). These findings are in line with previously reported sex-differences to specific antimigraine drugs, such as lack of inhibition of the spontaneous activity of trigeminocervical neurons by a CGRP-mAb in female rats [37], and higher headache recurrence rates in women after triptans [38]. Although (cycling) sex-hormones are thought to play a dominant role in modulating sex-based differences in pharmacokinetics and pharmacodynamics (reviewed in [39]), our study was not designed nor powered to study treatment differences across the estrous cycle. Clearly, additional studies are highly needed to address the exact neurobiological underpinnings behind these responses.

Lastly, we demonstrated that antagonism of the AMY_1_ receptor with AC 187, significantly attenuated pramlintide-induced delayed activation and sensitization of central trigeminocervical neurons in females (Figure 3 and Figure 4). Interestingly, innocuous somatosensory-evoked trigeminocervical neuronal responses were not inhibited in females with falling estrogen levels, whereas ongoing spontaneous and dural-evoked firing rates of trigeminovascular neurons were attenuated but not abolished. There are multiple explanations behind the incomplete blockage of the above responses. Firstly, activation of AMY_1_ receptors on C-fibers could release CGRP, which then diffuses and activates CGRP receptors on Aδ-fibers and possibly AMY_1_ receptors on C-fibers. However, this seems unlikely as AC 187 pretreatment fully blocked pramlintide-induced activation of Aδ-fibers. Secondly, pramlintide-induced responses could be mediated by activation of additional amylin responsive receptors, such as AMY_3_ receptors (see [20]), therefore AC 187 pretreatment would not be able to block these responses. However, this is also improbable as AC 187 has also affinity for AMY_3_ receptors [21], whereas other amylin receptors were not found in trigeminal nociceptors [40] and the low dose of pramlintide used is unlikely to activate additional receptors. Finally, if there is a higher expression of AMY_1_ receptors in females with falling estrogen (currently unknown but described with CGRP [3]), higher doses of AC 187 may be required to fully abolish pramlintide-induced responses. We did not test higher doses of AC 187 as the initial dose chosen was able to significantly decrease trigeminal nociception (Figure 2). Consequently, additional studies are required not only to evaluate whether AMY_1_ receptor expression varies between males and females, and females across the different phases of the estrous cycle, but also to dissect the neurobiological mechanisms that promote higher rates of sensitization in females, particularly during falling estrogen levels.

In conclusion, our results indicate that AMY_1_ receptor signaling modulates trigeminal nociception and these effects are more pronounced in females with falling estrogen levels. Therefore, blockage of these receptors may represent a potential drug target for the treatment of migraine.

## Key findings

Pramlintide, a selective AMY_1_ receptor agonist, increases ongoing spontaneous activity and dural-evoked responses only in females with falling estrogen levels.

AC 187, a selective AMY_1_ receptor antagonist, decreases dural-evoked responses only in males and ongoing spontaneous activity in males and females.

Pretreatment with AC 187 prevented pramlintide-induced increases in spontaneous activity and dural-evoked responses in females with falling estrogen levels.

